## Supplementary Figures for "The 3D spatial constraint on 6.1 million amino acid sites in the human proteome"

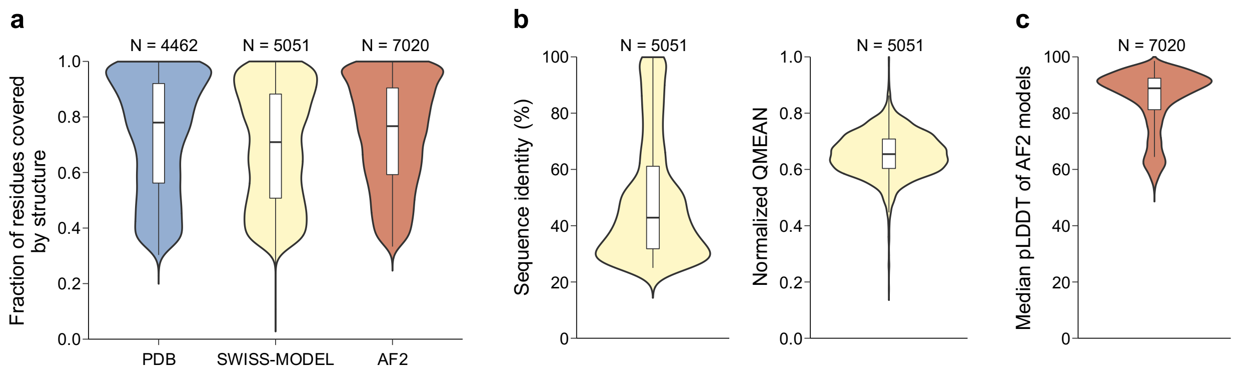


**Fig S1. Coverage and quality of protein 3D structures used in COSMIS. a**) Distribution of the fraction of residues covered by 3D structures for all proteins in the human reference proteome for which COSMIS scores are computed. Distributions are plotted separately for each of the three protein 3D structure sources. Median coverage of structures from PDB, SWISS-MODEL, and AF2 are 78.0%, 70.9%, and 76.7%, respectively. **b**) Left: Distribution of sequence identity of SWISS-MODEL homology models (median = 42.9%); Right: Distribution of normalized QMEAN scores in the range [0, 1] of SWISS-MODEL homology models (median = 0.654). **c**) For each AlphFold2 (AF2) protein 3D structure model used in COSMIS, we computed the median pLDDT score in the range [0, 100] of the residues for which COSMIS scores were calculated. The plot shows the distribution of median pLDDT scores of 7,020 AF2 models (median = 88.8).

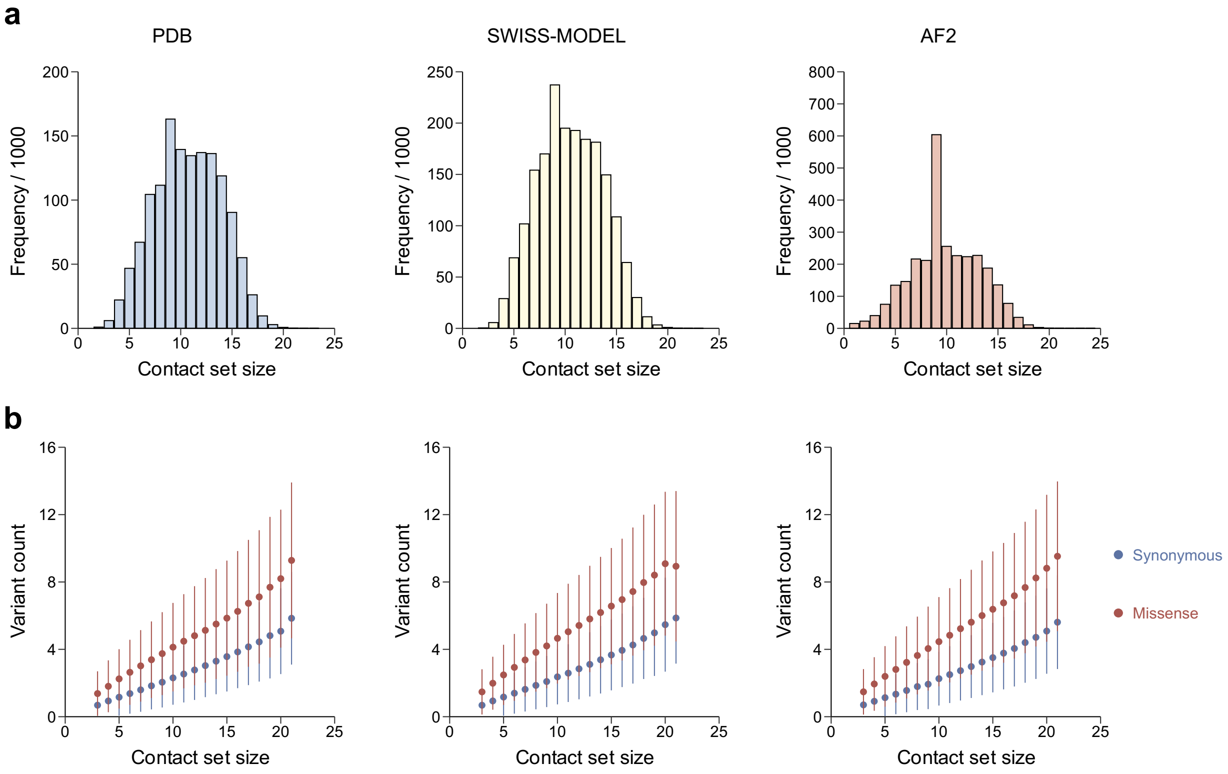


**Fig S2. Statistics of contact sets computed based on structures from PDB, SWISS-MODEL, and AF2, respectively. a**) Distributions of the size of contact set (number of amino acid sites in the contact set). The overall distributions are similar across difference protein 3D structure sources. **b**) Distribution of synonymous and missense variant counts across different sizes of contact set. Again, the overall distributions are similar regardless of the source of protein 3D structures.

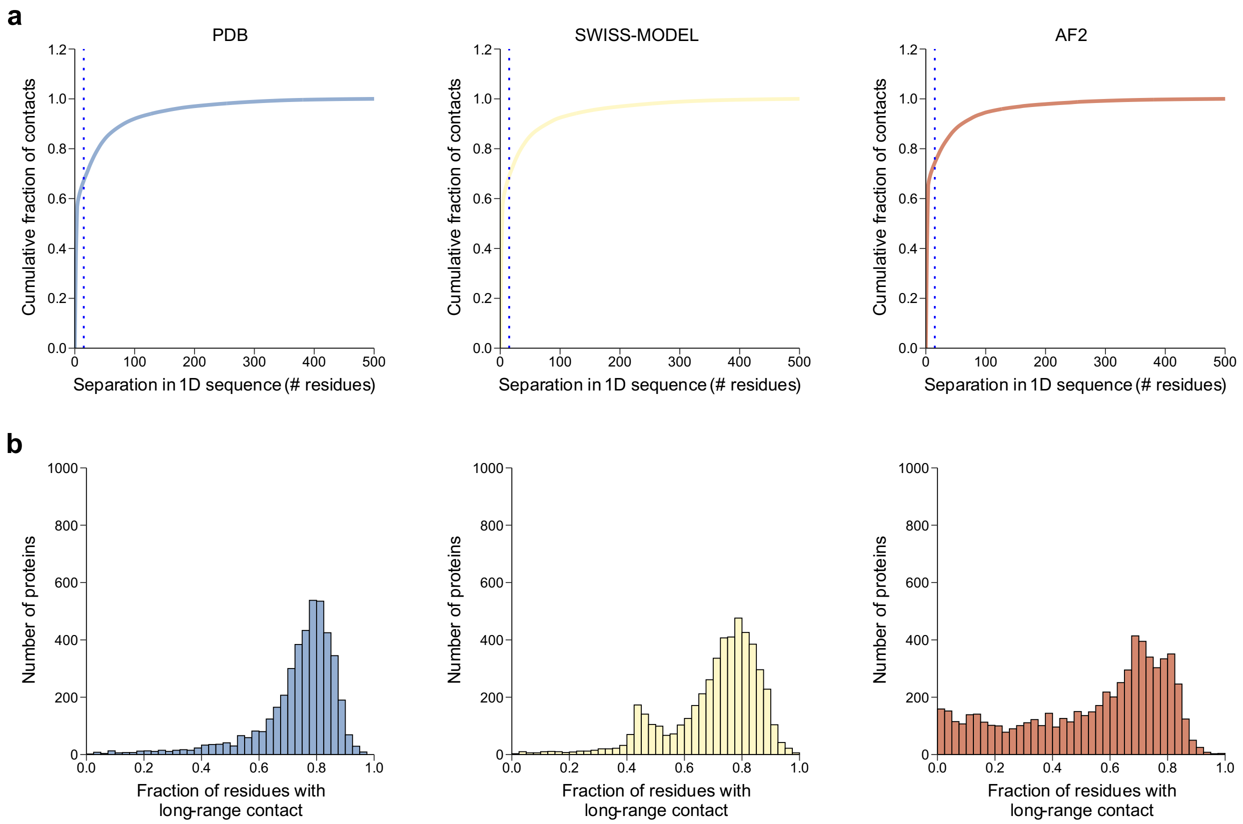


**Fig S3. Statistics on long-range contacts observed in protein 3D structures from the PDB, SWISS-MODEL, and AF2 databases. a**) Cumulative distribution of the sequence separation (number of residues apart in sequence) of all 3D contacts observed in structures from the three protein structure databases, respectively. Blue dotted lines correspond to a separation of 15 residues in 1D sequence. **b**) Distributions of per-protein fraction of residues that make at least one long-range contact.

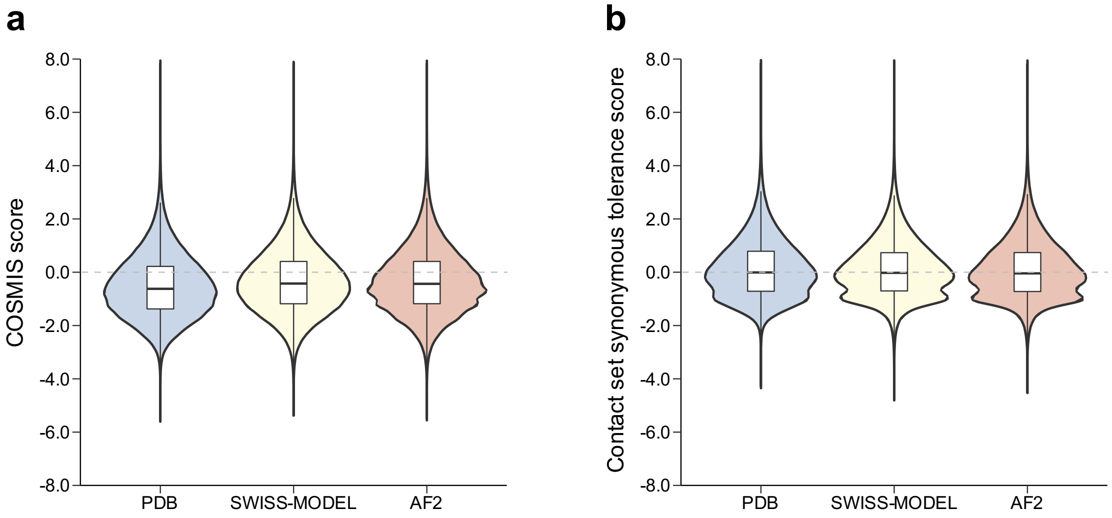


**Fig S4. Distributions of COSMIS (COntact Set MISsense tolerance) and contact set synonymous tolerance scores by sources of protein 3D structures. a**) Distributions of COSMIS scores computed based on difference sources of protein 3D structures. In general, proteins with experimentally determined structures in the PDB have a significantly lower COSMIS score than those that only have computationally predicted structures in SWISS-MODEL or AF2 databases (median -0.62 vs. -0.42 and 0.44, respectively, $p < 2.2 \times 10^{-308}$, two-sided Mann-Whitney U test). **b**) Distributions of contact set synonymous tolerance score computed based on difference sources of protein 3D structures. The scores are centered at the expected score under neutrality (i.e., median 0) regardless of the sources of protein 3D structures, consistent with the hypothesis that synonymous variants are not subject to 3D spatial constraint in protein structures.

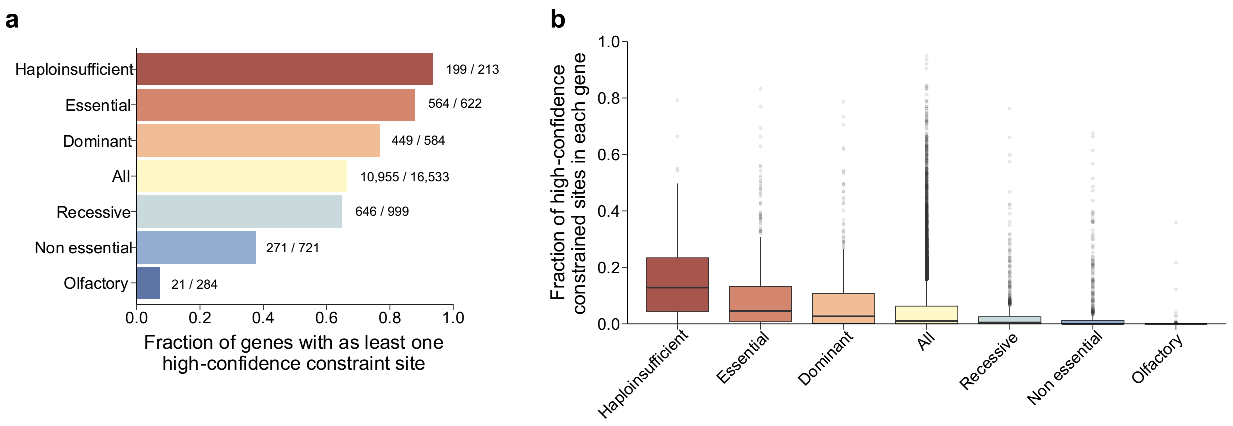


**Fig S5. Abundance of high-confidence constrained sites in genes with different levels of functional constraint. a**) Fraction of genes with at least one high-confidence constrained site in each of the six categories of genes. **b**) Distribution of per-gene fraction of high-confidence constrained amino acid sites of each gene category. In general, high-confidence constrained sites are more common in genes with essential functions and disease associations compared to genes with functions less essential to health and fitness.

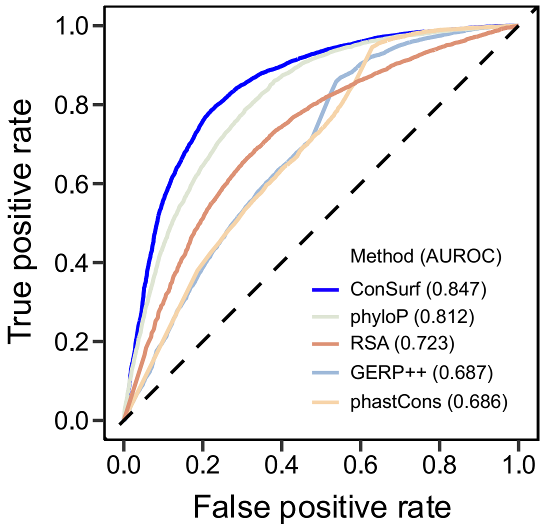


**Fig S6. Performance of phylogenetic conservation metrics in predicting variant pathogenicity.** The evaluation was performed on a total of 8,062 benign and 7,256 pathogenic missense variants from ClinVar for which all scores can be computed (Supplementary Table 6). We also evaluated the performance of relative solvent accessibility (RSA) in addition to four phylogenetic conservation metrics (GERP++, phyloP, phastCons, and ConSurf). Here, the best-performing metric is ConSurf. Note that in contrast to ConSurf, GERP++, phyloP, and phastCons quantify constraint on nucleotide sequence.

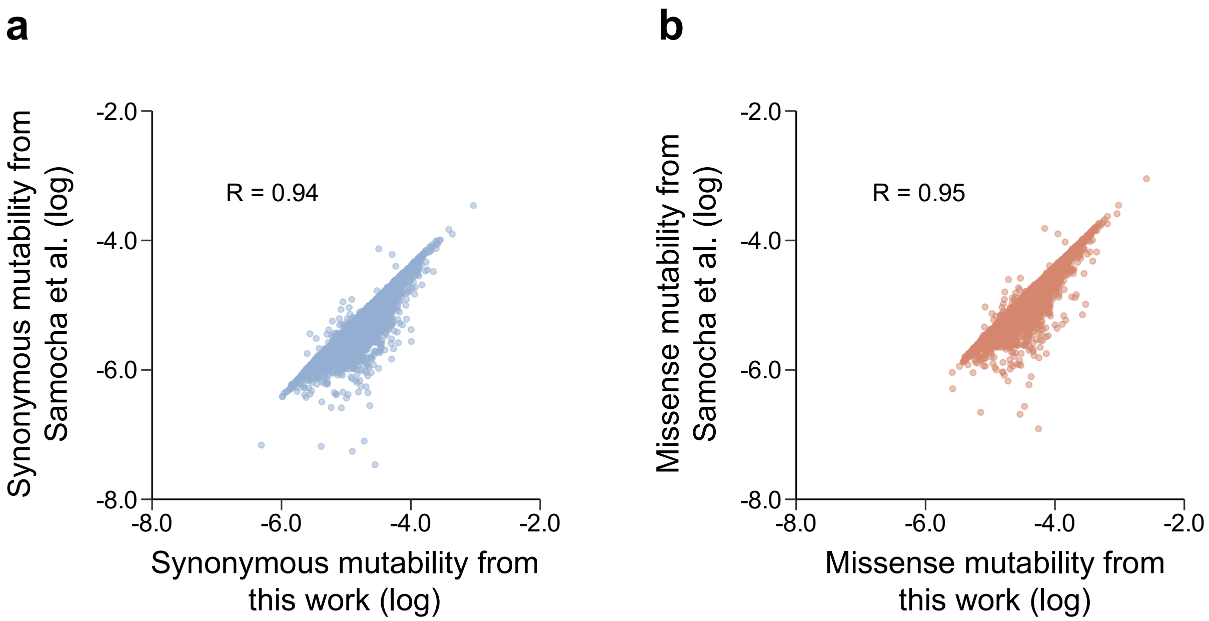


**Fig S7. Agreement between estimates of per-protein total synonymous and missense mutability used in COSMIS and previous work.** The scatter plots and Pearson's R are both based on the total synonymous and missense mutability of 14,756 Ensembl canonical transcripts with estimates from Samocha et al. 2014.

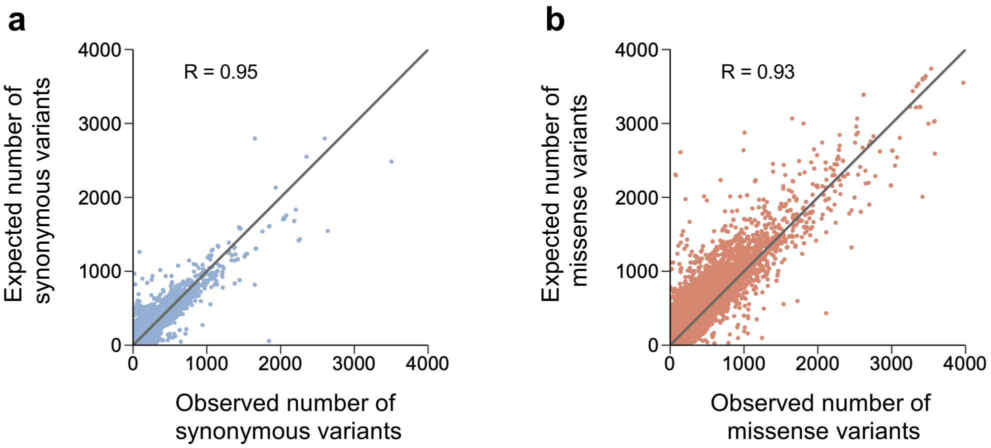


**Fig S8. Agreement between per-protein total number of observed and expected synonymous and missense variants.** The total expected number of variants per protein was computed based on the relation between synonymous variant count and synonymous mutability (from the fitted linear regression), i.e., $\hat{y} = 6.42 \times 10^{-6} \times \mu - 0.18$, where $\mu$ is per-protein total synonymous or missense mutability.

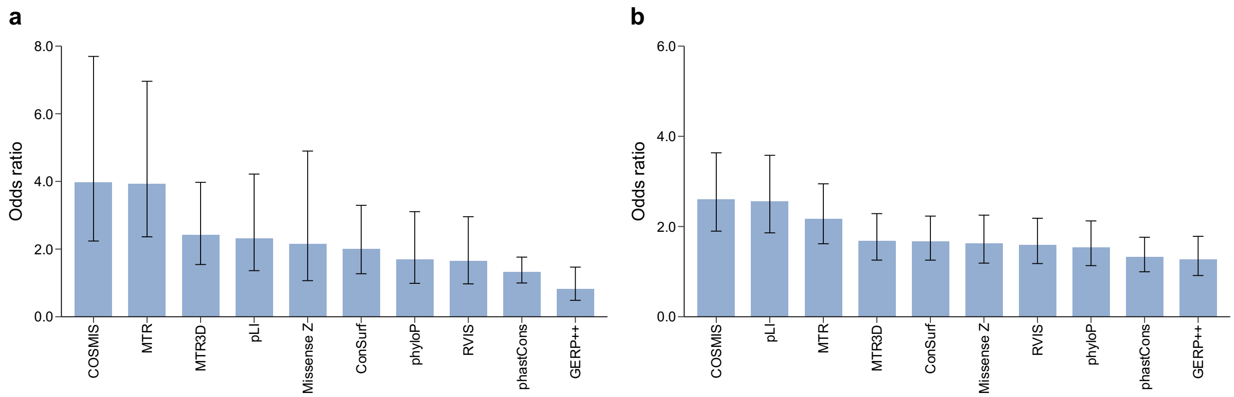


**Fig S9. Case variant enrichment analysis for intra- and inter-species constraint metrics at additional thresholds. a**) Odds ratios (ORs) of evaluated constraint metrics at the 5$^{th}$ percentile most constrained sites. COSMIS has the highest enrichment for cases (OR 4.0, 95% confidence interval [2.2, 7.7]). **b**) ORs of evaluated constraint metrics at the 20$^{th}$ percentile most constrained sites. While the ORs at 20$^{th}$ percentile threshold are generally lower than at 5$^{th}$ and 10$^{th}$ percentile thresholds, COSMIS still has the highest enrichment for cases (OR 2.6, 95% confidence interval [1.9, 3.6]) compared to all other evaluated metrics. Error bars are 95% confidence intervals of ORs.

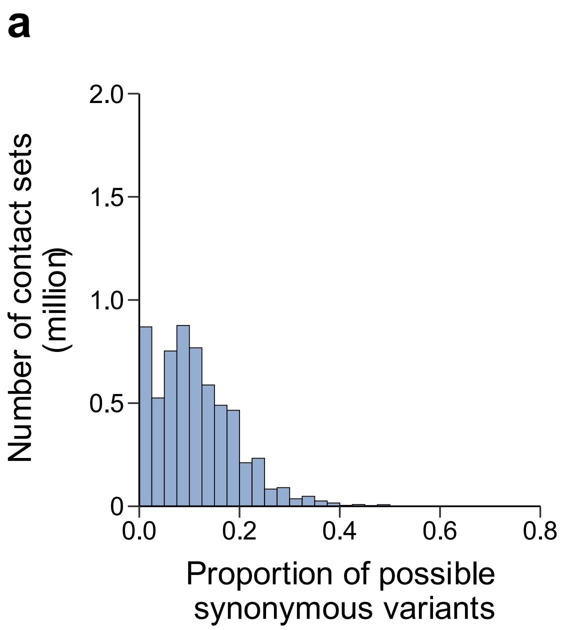


**Fig S10. Proportion of all possible a) synonymous and b) missense variants observed in each contact set.** On average, 10.3% and 6.3% of all possible synonymous and missense variants in a contact set are observed in gnomAD, respectively.
